# *In Silico* Characterization of Intracellular Localization Signals and Structural Features of Mosquito Densovirus (MDV) Viral Proteins

**DOI:** 10.1101/2023.12.13.571551

**Authors:** Tapan K Barik, Surya N Swain, Sushil Kumar Sahu, Usha R Acharya, Hillery C. Metz, Jason L Rasgon

## Abstract

As entomopathogenic viruses, mosquito densoviruses (MDVs) are widely studied for their potential as biocontrol agents and molecular laboratory tools for mosquito manipulation. The nucleus of the mosquito cell is the site for MDV genome replication and capsid assembly, however the nuclear localization signals (NLSs) and nuclear export signals (NES) for MDV proteins have not yet been identified. We carried out an *in silico* analysis to identify putative NLSs and NESs in the viral proteins of densoviruses that infect diverse mosquito genera (*Aedes, Anopheles*, and *Culex*) and identified putative phosphorylation and glycosylation sites on these proteins. These analyses lead to a more comprehensive understanding of how MDVs are transported into and out of the nucleus and lay the foundation for the potential use of densoviruses in mosquito control and basic research.

**Data summary:** Data used in this article were obtained from the GenBank database using accession numbers AYH52680, AYH52678, AYH52679, ABX83665, ABX83663, ABX83664, ABU95013, ABU95011, ABU95012, and AXQ04861.

**Impact statement:** Mosquito densoviruses (MDVs) are of interest as mosquito biocontrol agents and as laboratory research tools. The trafficking of viral proteins to the cell nucleus is a critical step in viral replication. We used *in silico* approaches to identify putative nuclear localization signals and nuclear export signals for MDVs that infect the three major genera of pathogen-transmitting mosquitoes (*Aedes, Anopheles*, and *Culex*). These analyses lead a more comprehensive understanding of how MDVs are transported into and out of the nucleus and lay the foundation for the potential use of densoviruses in mosquito control and basic research.

## INTRODUCTION

Densoviruses (DNVs) are pathogenic viruses in the family *Parvoviridae* that infect many invertebrate species (1-2). Mosquito densoviruses (MDVs) are mosquito-infecting, autonomous, non-enveloped, single-stranded viruses with a genome length of 4 to 6 kb. Though some are benign, MDV infections can be harmful to mosquitoes, adversely affecting multiple organs and potentially leading to larval deformation and death (3). Despite often being pathogenic, MDV infections are common in natural populations (3) and may spread horizontally between larvae in aquatic breeding sites (3-4), venereally between mating partners (5), and vertically from parent to offspring (3). Due to their pathogenicity, ability to spread in wild populations, and other advantageous properties such narrow host range, MDVs are considered to have potential as biological control agents and laboratory tools for mosquito manipulation (3-4).

The linear, single-stranded DNA genomes of DNVs encode proteins including the virion non-structural proteins (VPs) and virion capsid proteins (VCPs) (6-9). The number of capsid proteins varies among different genera of DNVs (10-12), but they all share common C-terminal and N-terminal regions and are associated with wide range of cellular functions including recognition of receptors for binding, delivery of the viral genetic elements to the nucleus, export of virion progeny during the late endosome period, and spread to other tissues (13). Intracellular trafficking of these viral proteins may contribute to successful viral replication and capsid assembly (14-15). Viruses often possess both localization and export signals that guide where they are trafficked in cells via energy-dependent protein transport (16-17).

Densovirus particles accumulate in the nuclei of the insect cells they infect (4), suggesting DNV proteins are trafficked into (and potentially later out of) the nucleus; a process typically mediated by peptide transport signals. Nuclear localization signals (NLSs) target proteins to the nucleus. They generally consist of a short polypeptide chain of positively charged amino acids and are classified as either classical or non-classical. Classical NLSs fall into two categories: monopartite and bipartite. Monopartite NLSs are short polypeptide chains of five to seven basic amino acids while the bipartite NLSs are composed of two clusters of basic amino acids separated by a linker of 10–11 amino acids. Apart from the monopartite and bipartite signals, the proteins involved in nuclear transport are also tagged with numerous localization signals such as proline-tyrosine and arginine-rich NLSs that do not follow the consensus pattern of classical NLS, referred to as “non-classical” (18). The transfer of functional proteins out of the nucleus to the cytoplasm through the nuclear pore complex is mediated by nuclear export signals (NES). A nuclear export signal (NES) generally consists of four hydrophobic amino acid residues according to Φ1-X2,3-Φ2-X2,3-Φ3-X-Φ4, where Φ represents Leu, Val, Ile, Phe, or Met, and X can be any amino acid (19). *In silico* approaches can be used to predict NLSs and NES regions within a protein sequence (20).

We searched for the presence of putative intracellular trafficking signals from mosquito densovirus protein sequences. We carried out a heuristic search of DNVs infecting the three mosquito genera that most strongly impact global public health: *Aedes, Anopheles*, and *Culex*. We identified functional protein domains and predicted sites targeted for post-translational modifications. These analyses will lead to a more comprehensive understanding of MDVs capsid protein function and lay the groundwork for potential DNV modifications for applied use.

## METHODS

### Sequence data

From the literature we identified MDVs that infect *Aedes, Culex*, and *Anopheles* mosquitoes: *Aedes albopictus* densovirus (AalDNV), *Anopheles gambiae* densovirus (AgDNV), and *Culex pipiens* densovirus (CpDNV). The translated amino acid sequences for the VPs/VCPs of these viruses (accession numbers AYH52680, AYH52678, AYH52679, ABX83665, ABX83663, ABX83664, ABU95013, ABU95011, ABU95012, and AXQ04861) were retrieved from GenBank (National Center for Biotechnology Information, available at https://www.ncbi.nlm.nih.gov). MDVs VP/VCP ontology annotations and predicted functional classifications are given in Table 1.

**Table 1:** Summary of predicted functional classifications of MDV capsid proteins.

| Biological Process |  | Cellular Component |  |
| --- | --- | --- | --- |
| GO ID | Reliability Score (%) | GO ID | Reliability Score (%) |
| Interspecies Interaction between Organisms; GO:0044419 | 60 | Virion; GO:0019012 | 63 |
| Viral life cycle; GO:0019058 | 60 | Virion part; GO:0044423 | 43 |
| Interaction with host; GO:0051701 | 60 | Viral capsid; GO:0019028 | 43 |
| Symbiosis, encompassing mutualism through parasitism; GO:0044403 | 60 | T=1 Icosahedral viral capsid; GO:0039615 | 35 |
| Virion attachment to host cell; GO:0019062 | 47 | Icosahedral viral capsid; GO:0019030 | 35 |
| Viral process; GO:0016032 | 47 |  |  |
| Viral entry into host cell; GO:0046718 | 46 |  |  |
| Entry into other organism involved in symbiotic interaction; GO:0051828 | 46 |  |  |
| Movement in host environment; GO:0052126 | 46 |  |  |
| Locomotion; GO:0040011 | 46 |  |  |

### Functional Prediction of Viral Proteins

To identify putative functions associated with MDV capsid proteins, we carried out two sequence-based analyses: Protein Function Prediction (PFP) (21-22) and Support vector machine (SVM)-Prot (23). PFP is an algorithm to predict the most probable protein function (in Gene Ontology [GO] terms) of a submitted query sequence. We used the PFP server PSI-BLAST (blastpgp version: 2.2.10) to predict the most probable GO biological processes and cellular component functions. PFP uses a scoring scheme (‘reliability’) to rank GO annotations based on frequency and degree of similarity of the originating sequence to the query sequence (21-22). SVM-Prot classifies proteins into functional families through a machine learning method from primary query protein sequences. We used the SVM-Prot web server (23) to predict potential specific functions based on physicochemical properties such as amino acid composition, charge, polarity, polarizability, molecular weight, solubility, surface tension, hydrophobicity, Van der Waals force, secondary structure, solvent accessibility, and number of hydrogen bond donors and side chain acceptors.

### Prediction of Subcellular Localization of Viral proteins

We used the pLOC-mVirusweb-server (24) to predict subcellular protein locations in terms of optimal GO information by combining with the composition of pseudo amino acids (PseAA) in the protein structure.

### Search for Intracellular Localization Signals

We used cNLS Mapper (25), NLStradamus (26), NucPred (27), and SeqNLS (28) servers to make Nuclear Localization Signal (NLS) predictions, and NetNES (29), NESFinder 0.2 (30) and LocNES (31) for Nuclear Export Signal (NES) predictions. The region(s) of the polypeptide chain that exceeded the cutoff value limit were considered to be involved in nuclear signaling.

### Prediction of Transmembrane helices and their topology

It is unusual to find transmembrane helices in the proteins of non-enveloped viruses, but they do occur. For example, a capsid protein of the Bluetongue virus (VP5) gained a transmembrane anchor region via fusion (32). We therefore searched for the presence of transmembrane helices in MDV capsid proteins using the TMHMM server 2.0 (33). TMHMM is a support vector machine (SVM) based web-server tool that works in a hierarchical framework to predict transmembrane helices and with their topology (32). Polypeptide secondary structure was predicted using the Raptor X server (35). To make accurate predictions, RaptorX uses a combination of three algorithms: deep convolutional neural fields (DCNF), deep convolutional neural network (DCNN), and conditional random fields (CRF).

### Viral protein 3D model prediction

The SWISS-Model Workspace (34) was used to homology model the structure of the *Aedes albopictus* densovirus capsid protein using the capsid protein of *Galleria mellonella* densovirus (1DNV) as the template sequence.

### Analysis of Post-translational modification sites

We used PROSITE (36) to identify potential glycosylation and phosphorylation sites in the MDV (AgDNV) capsid protein sequence.

## RESULTS

### Densovirus viral proteins predicted to have essential functions

Protein Function Prediction (PFP) analysis associated MDV viral proteins with multiple functions including viral life cycle, interaction with host symbiosis and mutualism, virion attachment to host cell, and viral entry into host cell (Table 1).

### MDV viral proteins contain both NLS and NES nuclear transport signals

We identified intracellular NLS and NES trafficking signals in all MDV proteins. Identified motifs were classical monopartite NLSs and their arrangement agreed with the consensus sequence of K(K/R)X(K/R) and K/R)(K/R)X10–12(K/R)3/5 respectively, where X represents any amino acid and (K/R)3/5 indicates the presence of three out of five consecutive basic amino acids (37) (Table 2). The NLSs predicted by cNLS Mapper were high-score monopartite motifs in the N-terminal end of VP/VCP proteins of MDVs. We also found that all investigated viral capsid protein sequences contained nuclear export signals (NESs) (Table 3).

**Table 2:**
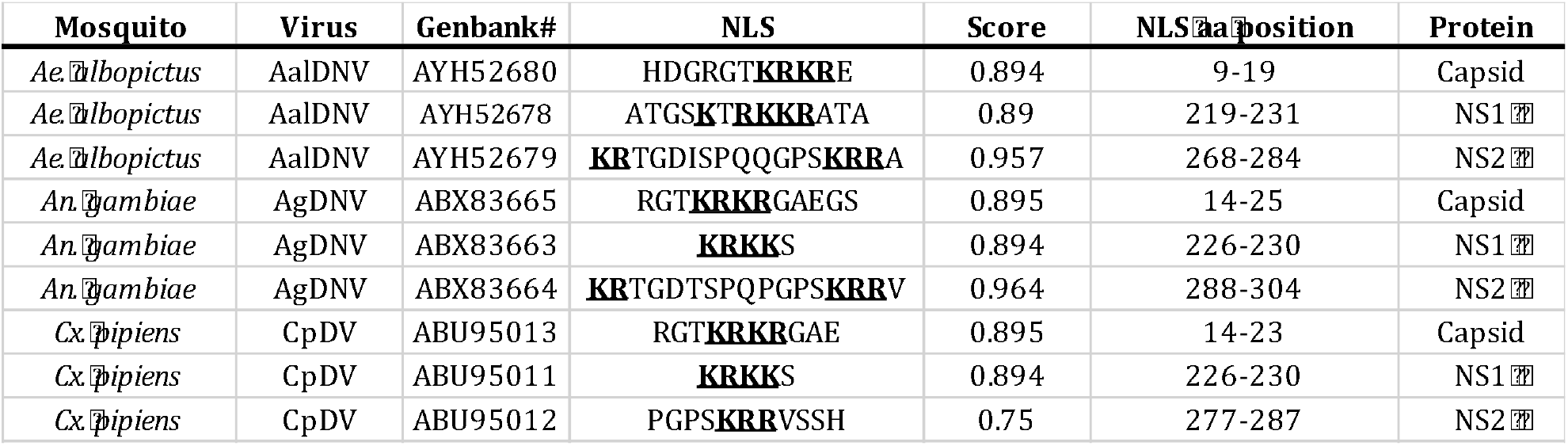
Predicted NLSs in the proteins of MDVs infecting 3 mosquito genera. Bold underlined amino acid sequences are monopartite NLSs with the consensus K(K/R)X(K/R) (X = any amino acid residue).

**Table 3:** Predicted NESs in the capsid proteins of MDVs infecting 3 mosquito genera. Leucine residues are bolded and underlined.

| **Mosquito** | **Virus** | **GenBank#** | **NES** | **Score** | **aa position** | **Protein** |
| --- | --- | --- | --- | --- | --- | --- |
| *Ae. albopictus* | AaDNV | AYH52680 | NQSINIMRAL<u>**NAVAL**</u> | 0.19 | (103-117) | Capsid |
| *An. gambiae* | AgDNV | ABX83665 | SSTNMEYMQRQ<u>**VLEL**</u> | 0.27 | (323-337) | Capsid |
| *Cx. pipiens* | CuDNV | AXQ04861 | SSTNIEYMHRQ<u>**VLEL**</u> | 0.2 | (323-337) | Capsid |

### MDV capsid proteins are predicted to be monomers with a primarily coiled secondary structure supported by stabilizing interactions

We predicted secondary structure of MSV capsid proteins and found coil residues were more common than strand and helix-residues. Specifically, capsid proteins had >60% coil residues (*Aedes albopictus* DNV capsid shown as an example in Figure 1; the percentage of strand and helix-residues was similar for all MDV capsid proteins examined). We selected the capsid protein of *Galleria mellonella* densovirus 1 as template for 3D model prediction (30.08% identity to the query; *Aedes albopictus* DNV capsid). The oligomeric state of the predicted model was a monomer (Figure 2).

**Figure 1:**
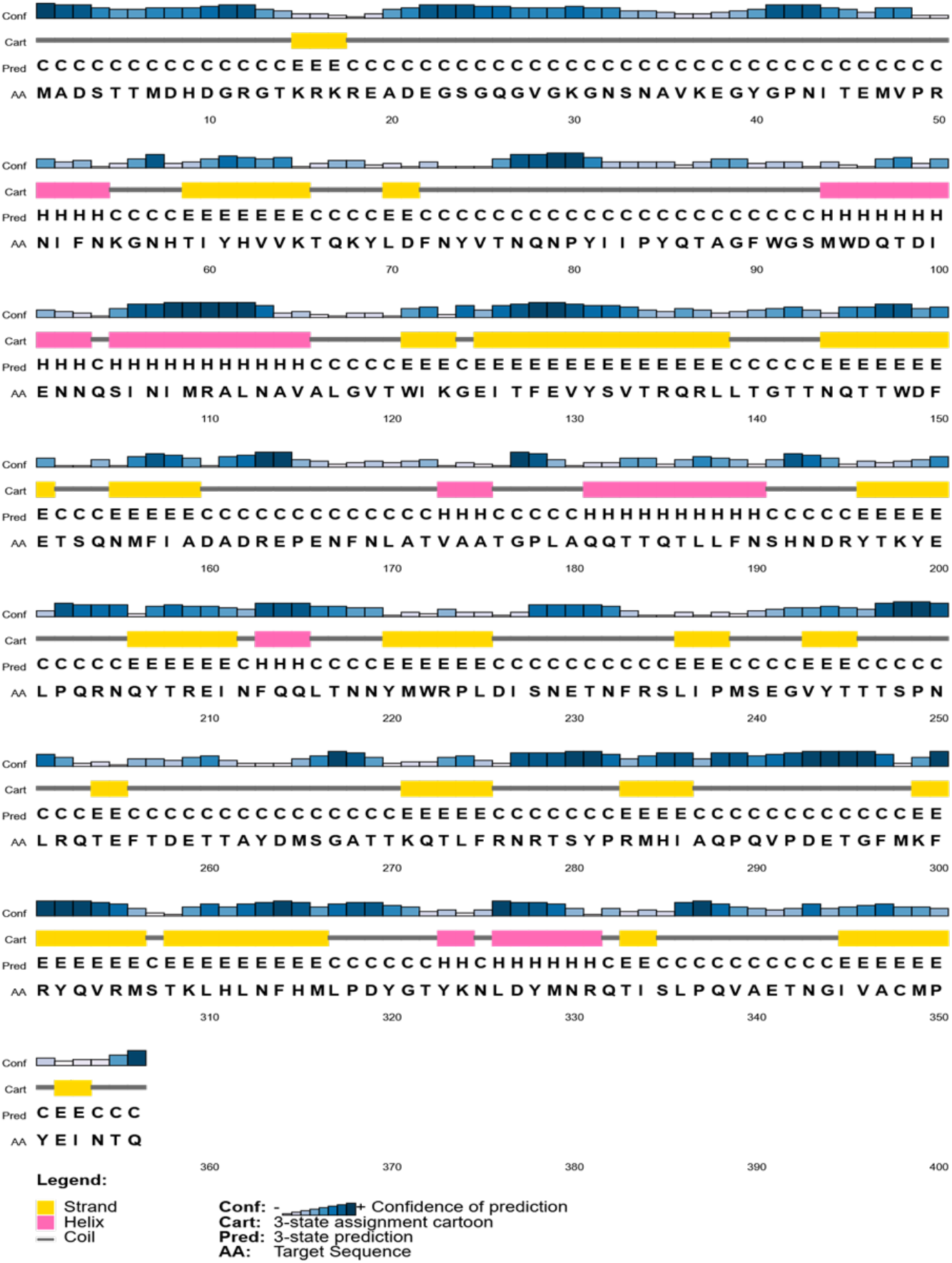
Secondary structure analysis of MDV capsid protein (*Aedes albopictus* DNV used as type sequence).

**Figure 2:**
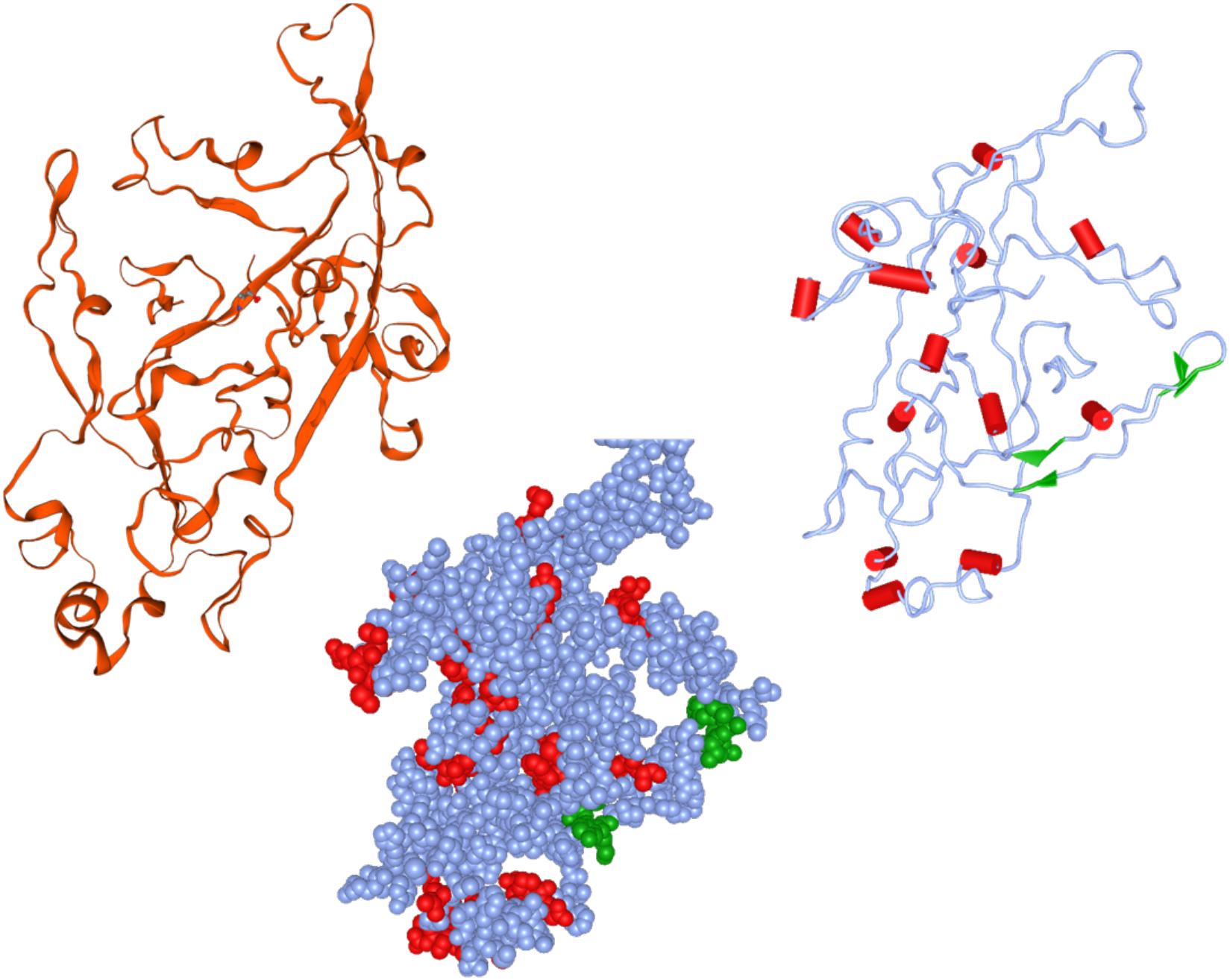
Three-dimensional homology model of the MDV capsid protein (*Aedes albopictus* DNV). (a) Ribbon model, (b) Cylinder and plate model, and (c) Sphere model. Strand residues are highlighted in green, helix residues are highlighted in red.

### MDV VP protein function may rely on phosphorylation

PROSITE analysis indicated that the MDV (AgDNV) capsid protein possesses abundant putative phosphorylation and glycosylation sites. The serine and threonine residues were predicted to be targeted sites for phosphorylation (Table 4).

**Table 4:** MDV capsid predicted consensus phosphorylation and glycosylation sites for the AgDNV capsid protein (CK2: Casein kinase II; PKC: Protein kinase C).

| CK2-PHOSPHORYLATION SITES |  |  |  |
| --- | --- | --- | --- |
| aa position | aa sequence | Modified residue | Phosphorylated residue |
| 5-8 | TSMD | 5 | Phosphothreonine |
| 36-39 | SVKE | 36 | Phosphoserine |
| 94-97 | SMWD | 94 | Phosphoserine |
| 147-150 | TTWD | 147 | Phosphothreonine |
| 198-201 | TKYE | 198 | Phosphothreonine |
| 325-328 | TNME | 325 | Phosphothreonine |
| PKC-PHOSPHORYLATION SITES |  |  |  |
| 16-18 | TKR | 16 | Phosphothreonine |
| 36-38 | SVK | 36 | Phosphoserine |
| 64-66 | TVK | 64 | Phosphothreonine |
| 267-269 | SGK | 267 | Phosphoserine |
| 308-310 | STK | 308 | Phosphoserine |
| 341-343 | TAR | 341 | Phosphothreonine |
| N-GLYCOSYLATION SITES |  |  |  |
| 44-47 | NMTE | - | N-Linked (GlcNAc) Asparagine |
| 58-61 | NHTV | - | N-Linked (GlcNAc) Asparagine |
| 104-107 | NNTI | - | N-Linked (GlcNAc) Asparagine |
| 145-148 | NQTT | - | N-Linked (GlcNAc) Asparagine |
| 278-281 | NRTS | - | N-Linked (GlcNAc) Asparagine |

## DISCUSSION

Densoviruses (DNVs) are some of the most important and intensively studied mosquito pathogenic viruses (38). DNVs characteristics such as environmental stability, narrow host range, and pathogenicity make them particularly suited to biocontrol applications (39-42).

Although intercellular trafficking is critical in the MDV life cycle (43), less is known regarding the involvement of specific intracellular NLSs and NES. To address this gap, we analyzed all MDV viral protein sequences from three medically important mosquito genera for the presence of intracellular trafficking signals *in silico* using bioinformatics tools and databases. We found that the viral proteins of MDVs have signal peptides with both nuclear localization and export functions.

The distribution of identified trafficking signals agrees with the biology of MDV. All three MDV proteins (capsid, NS1, and NS2) possess NLSs, as these viral proteins must be trafficked into the nucleus for viral replication (43). However, only the capsid proteins possessed a predicted NES, as the virion must exit the nucleus to infect other cells or to be released into the environment. MDVs often form paracrystalline arrays of virions in the cytoplasm of infected cells without the destruction of the nuclear envelope (44), and the presence of an NES could explain the export of newly formed virions from the nucleus to cytoplasm through the nuclear pore complex, thereby preserving the nuclear envelope. In sum, the presence of NLS and NES sequences in densovirus capsid proteins likely play a crucial role viral life cycle events such as budding and genome assembly.

It was found that nuclear localization signal peptides are largely conserved across three diverse mosquito genera. This level of conservation is significant, as *Anopheles* are believed to have split from other *Culicidae* in the Triassic, over 200 million years ago (45). These *in silico* findings may point to the specific motifs that control the nuclear translocation of these peptides. Our predictions of the secondary structure along with some crucial features such as functional domains, phosphorylation and glycosylation sites, and 3D structure provide novel structural insight of these MDVs proteins.

In conclusion, the presence of both NLS and NES in the MDV capsid protein suggests protein translocation to the nucleus for virus capsid assembly and subsequent release of the mature virus to the cytoplasm, whereafter it goes on to infect neighboring cells. The analyses reported here will lead to a more comprehensive understanding of MDVs capsid protein function, and lay the groundwork for targeted manipulations of DNVs that could be used for mosquito control and other applied uses.

## AUTHOR CONTRIBUTIONS

Conceptualization: TKB, JLR. Data Curation: TKB; Formal Analysis: TKB, SNS, SKS, URA; Funding acquisition: TKB, JLR; Investigation: TKB, SNS, SKS, URA; Methodology: TKB, JLR; Project administration: TKB; Resources: N/A; Software: N/A. Supervision: TKB; Validation: TKB; Writing – Original draft: TKB, HCM, JLR; Writing – Review and editing: TKB, JLR, HCM.

